# Self-multimerization of mRNA LNP-derived antigen improves antibody responses

**DOI:** 10.1101/2025.09.09.675153

**Authors:** Cody A. Despins, James Round, Lisa Dreolini, Tracy S. Lee, Scott D. Brown, Robert A. Holt

## Abstract

**Background:** mRNA LNP technology is now being widely applied as a highly effective vaccine platform. Antigen multimerization is a well-established approach to enhance antibody titers and protective efficacy of several protein subunit vaccines. However, this approach has been less explored for mRNA LNP vaccines.

**Methods:** Here, within the context of mRNA LNP vaccination, we used mStrawberry (mSb) as a model antigen to conduct a comprehensive, head-to-head comparison of the ability of the foldon (3-mer), IMX313 (7-mer), and ferritin (24-mer) multimerization domains to enhance immunogenicity in mice.

**Results:** We compared multimerized antigen to monomeric secreted antigen and monomeric surface-displayed antigen and observed that the IMX313 domain efficiently multimerized mSb protein and enhanced anti-mSb antibody titers to a degree greater than surface-display, whereas the foldon and ferritin domains failed to multimerize or improve antibody levels.

**Conclusions:** Our results extend the observation of enhanced immunogenicity from antigen multimerization to mRNA LNP vaccines and indicate the 7-mer forming IMX313 multimerization domain may be an ideal candidate for multimer formation in the context of mRNA LNP vaccination.

## Introduction

Widespread use of mRNA LNP vaccines during the COVID-19 pandemic (mRNA-1273 and BNT162b2) demonstrated the mRNA LNP platform’s capacity for both rapid development (critical for pandemic response and updating for variant emergence) and high protective efficacy. Since its debut in this setting, mRNA LNP vaccine development has expanded to target many other pathogens. mRESVIA (mRNA-1345) demonstrated an 83.7% protection from RSV-associated lower respiratory tract disease (LRTD) with ≥2 signs/symptoms in older adults in a phase II/III clinical trial [1], and was approved for use in older adults (≥60 years of age) by the FDA in 2024, representing the first non-SARS-CoV-2 mRNA LNP vaccine approval. In 2025, mRESVIA was further approved by the FDA for adults (18-59 years of age) with increased risk for LRTD, and it is currently being evaluated for use in children, another population with high risk of severe outcomes from RSV, in clinical trials [2,3]. A multiantigen CMV mRNA LNP vaccine (mRNA-1647) is currently being evaluated in a phase III clinical trial following promising phase I and II results [4,5]. A seasonal influenza mRNA LNP vaccine demonstrated superior immunogenicity variables to conventional protein vaccine comparators in phase III clinical trials and is also being evaluated as a flu/COVID-19 combination vaccine (mRNA-1010 and mRNA-1083, respectively) [6]. Many mRNA LNP vaccines targeting other viruses, including those for VSV, HIV, Zika, mpox, rabies, and EBV, are in earlier stages of clinical trial evaluation [7]. Further, mRNA LNP vaccines for bacterial targets, such as *Borrelia* spp. (the causative agent of lyme disease), and parasites, such as *Plasmodium falciparum* (the causative agent of malaria), are also in clinical trials [7]. These examples highlight the extensive innovation in infectious disease vaccine development enabled by mRNA LNP technology.

In addition to infectious disease applications, the ability to rapidly develop and test mRNA LNP vaccines makes personalized neoantigen vaccines a more viable therapeutic option in the field of oncology. Once a patient’s tumor is sequenced and putative neoantigens are identified, neoantigen-encoding mRNA LNP vaccines can be formulated and delivered to boost anti-tumor immunity, often in combination with other immunotherapies. This strategy has recently shown promising results in clinical trials for pancreatic cancer [8] and melanoma [9]. Furthermore, several non-personalized cancer mRNA LNP vaccines encoding common cancer associated antigens are also being developed [7].

Overall, mRNA LNP technology has revolutionized modern vaccinology, and promises major reduction of disease burden in both infectious disease and oncology, as well as potential for further improvement and applications as the technology develops.

All vaccines rely on being sufficiently immunogenic to drive the expansion of target-specific populations to a degree such that there is reduction or prevention of disease upon subsequent exposure (for infectious disease applications) or to promote re-engagement of immunity with the tumor (for oncology applications). Thus, in both settings, generating sufficient immunogenicity is vital, and towards this goal, it is well established that vaccination using multimerized antigen can enhance antibody responses. There are two primary proposed mechanisms for this: firstly, increased antigen avidity through multimerization can enhance B cell activation [10], and secondly, soluble antigen of an appropriate size (10-100 nm) can transit to draining lymph nodes independent of antigen-presenting cells (APCs) [11–13], thereby increasing concentration of antigen and likelihood of interactions with reactive B cells.

Several domains are commonly used to achieve antigen multimerization. The foldon domain, derived from fibritin of bacteriophage T4 [14], mediates formation of a 3-mer and has demonstrated enhanced neutralization of pseudoviruses compared to non-foldon vaccine, and complete protection against lethal (10 x LD_50_) challenge in murine models for influenza [15]. Foldon has also been applied for SARS-CoV [16], MERS-CoV [17], SARS-CoV-2 [18], RSV [19], and JEV vaccines [20]. Similarly, the IMX313 domain, which is derived from the chicken C4bp oligomerization domain [21] and mediates formation of a 7-mer, has been shown to significantly increase IgG levels at low doses and improve parasite transmission blockage compared to monomeric protein in a Pfs25-targeted malaria vaccine [22]. Lastly, the ferritin multimerization domain, commonly derived from *H. pylori*, mediates the formation of a 24-mer. An RBD-encoding ferritin vaccine approach increased antigen uptake by macrophages and DCs, and increased antibody titers and proportions of reactive CD8^+^ and CD4^+^ T cells following vaccination, compared to monomeric RBD [23]. Ferritin has also been used in vaccines for influenza [24,25], EBV [26,27], HIV [28], Zika [29], and Lyme disease [30].

Multimerization using foldon, IMX313, ferritin and other domains has been well characterized and broadly applied for protein vaccines. Similar findings have been reported for pDNA vaccines [31,32] and viral vector vaccines [22]. However, evaluation of antigen multimerization in the context of mRNA LNP vaccines remains limited. Whether the mRNA LNP platform is fully amendable to antigen multimerization (i.e. whether the dynamics of mRNA LNP-derived expression allow antigens to self-multimerize at a meaningful rate within the cell), if there are important considerations compared to multimerization for protein vaccines and other vaccine types, and whether some multimerization domains may be better suited for mRNA LNP application, are important questions to address towards the development of more effective mRNA LNP vaccines.

In this study, we aimed to address this knowledge gap by evaluating multimerization in the setting of mRNA LNP vaccination. Using mSb as a representative antigen of interest, we tested 3 commonly used multimerization domains (foldon [3-mer], IMX313 [7-mer], and ferritin [24-mer]) and evaluated their capacity to multimerize, secrete from the cell, and enhance antibody responses induced by mRNA LNP vaccination.

## Results

### Construct design & antigen structure prediction

Plasmid DNA (pDNA) constructs for generating mRNA for encapsulation and transfection/vaccination were designed incorporating several features for enhanced antigen expression, immunogenicity and ease of use. These constructs have a T7 promoter followed by an AG initiator for use with TriLink CleanCap AG during *in vitro* transcription (IVT) reactions to generate 5’ Cap-1 (m7GpppN1mp) capped mRNA (which, as opposed to Cap-0 [m7GpppN1p], has an additional 2’-O-methylation at the nucleotide 1 position). Such Cap-1 capping has been shown to be essential in mimicking self mRNA of higher eukaryotes, which has 2’-O-methylation at the nucleotide 1 position, and thereby reduces type I interferon activation and translational inhibition typically induced by uncapped or Cap-0 mRNAs [33–35]. The construct is also designed such that the IVT transcript has a 5’ α-globin UTR and a eukaryotic Kozak sequence motif for efficient protein translation [36,37], and 2x 3’ β-globin untranslated regions (UTRs) for greater mRNA translation/stability [38]. mRNA CDS were codon optimized using GenSmart codon optimization to enhance translation efficiency. The poly-A tail is 120bp to promote translation/stability [38], bisected to reduce pDNA recombination [39] and directly incorporated into the plasmid sequence (rather than added following IVT via enzymatic polyadenylation, which results in inconsistent polyA lengths [38]). The construct was also designed for linearization prior to IVT via a type IIS restriction enzyme (SapI or BspQI), eliminating any 3’ overhang at the end of the polyA tail following linearization, which have been reported to inhibit mRNA translation/stability [38]. Lastly, within the pDNA construct, an upstream CMV promotor enables expression of the mRNA CDS and thereby allows for *in vitro* assessment of designs by pDNA transfection in mammalian cells, such as HEK293T/17 cells. In addition, generation of mRNA was also performed using methodology for enhanced antigen expression and immunogenicity – during in vitro transcription, N^1^-methylpseudouridine was incorporated to reduce TLR activation and associated decreases in cellular translation [40].

For the mRNA CDS sequence, four different designs were created: for intracellular antigen expression, for expression and secretion of antigen as a monomer, for expression and secretion of antigen as a multimer, and for expression and surface-display of antigen (**Figure 1**). Given that mRNA LNP-encoded antigens are expressed in the cytoplasm, the mRNA CDS design for intracellular expression contains the antigen of interest with no additional domains. To stimulate a more robust antibody response, the vaccine-encoded antigen should be accessible to the extracellular environment. Therefore, a murine-derived Igκ signal peptide was added to the N-terminus of the antigen of interest for the mRNA CDS for secretion of monomeric antigen, which targets the encoded protein to the ER for subsequent secretion in mammalian cells [41– 43]. Multimerization domains (foldon-based, IMX313-based, or ferritin-based) were added in the C-terminal position in combination with Igκ signal peptide to generate an mRNA CDS for secretion of multimeric antigen. Surface-display is a commonly used alternative approach to extracellularly expose antigen. Furthermore, surface-displayed RBD-encoding saRNA LNP vaccines have shown greater antibody titers compared to secreted monomeric RBD [44]. Therefore, designs for surface-display were also created for direct comparison to secreted multimeric antigen designs. For the mRNA CDS for surface-displayed designs, the Igκ signal peptide is included in combination with a C-terminal transmembrane domain, either B7 (including the cytoplasmic tail) or PDGFR-based. In combination with the ER-directing signal peptide Igκ, both B7 TM domain and PDGFR TM domains have been shown to lead to surface-display of the protein [45–47]. Lastly, a StrepTagII sequence was included at the C-terminal end of constructs. This enables immobilization of antigen for various downstream applications, such as ELISA (using streptavidin-coated plates) or purification. StrepTagII sequences were chosen as tags, rather than other commonly used tags, such as HA or Myc sequences, which are microbe-derived and therefore potentially highly immunogenic. The StrepTagII sequences were not included in the mRNA CDS designs for secretion of multimeric antigen, given concerns that it may interfere with multimerization. For simplicity, these various mRNA CDS are named according to their features from N-terminus to C-terminus (e.g. Igκ-mSb-TM(B7)-Strep) and are referred to as “intracellular”, “secreted monomeric”, “secreted multimeric”, or “surface-displayed” designs.

**Figure 1.**
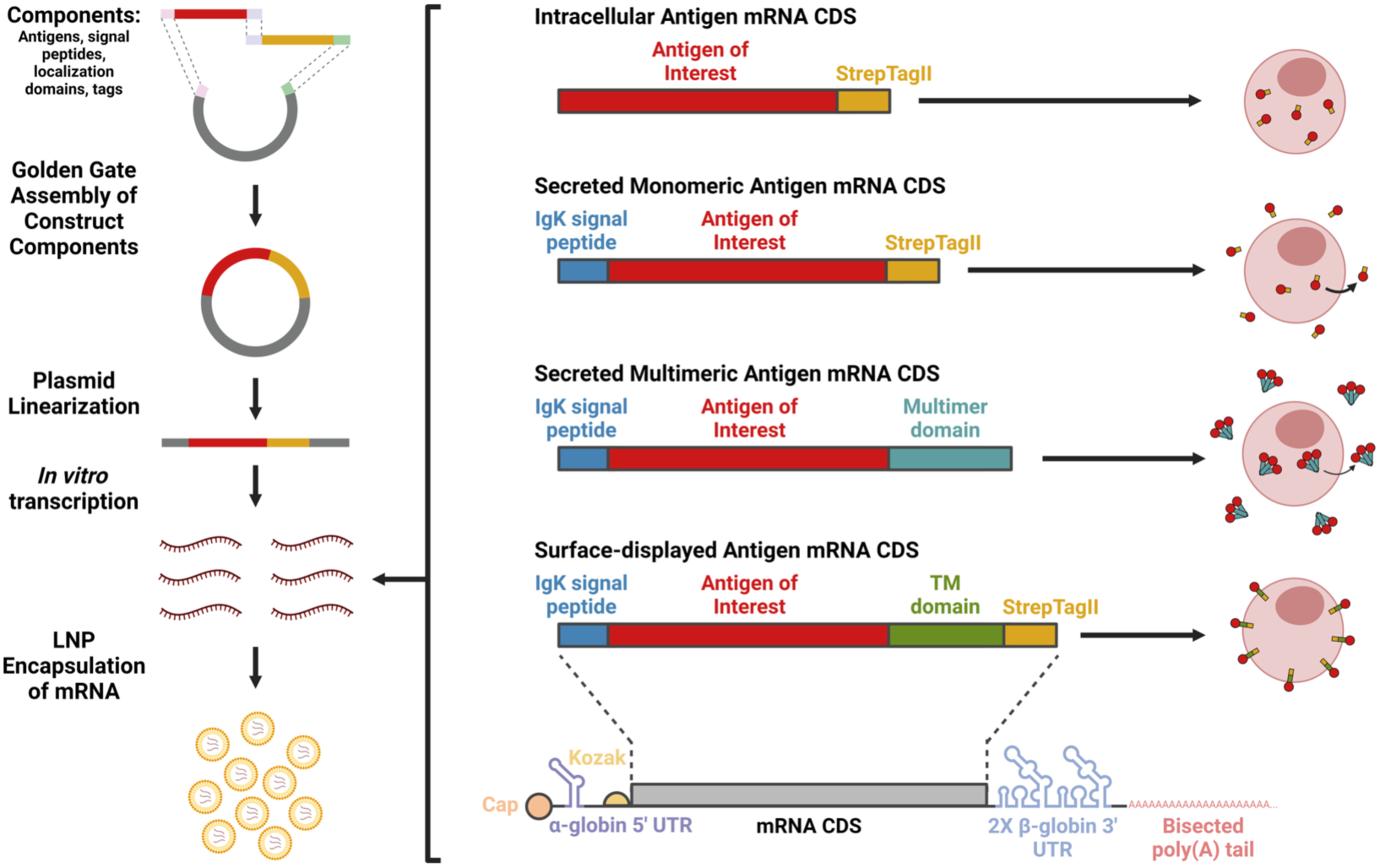
Overview of mRNA LNP vaccine production pipeline and design types. Construct design components such as antigen, signal peptides, domains for localization (such as transmembrane [TM] domains for surface-display) and tags are assembled into a pDNA construct using Golden Gate cloning methods. Plasmids are then isolated, linearized, and *in vitro* transcription is performed to generate messenger RNA (mRNA). mRNA is encapsulated in lipid nanoparticles (LNP) and evaluated *in vitro* via transfection and *in vivo* via intramuscular (IM) vaccination. Four primary mRNA CDS designs were used for: intracellular expression of antigen, expression and secretion of the antigen as a monomer, expression and secretion of the antigen as a multimer (3-mer foldon-based multimer shown), and expression and surface-display of the antigen. Figure was made using BioRender.com.

We firstly performed structural predictions of the different secreted multimeric design types using AlphaFold2 [48] to determine whether the structures were amendable to expected multimer formation (**Figure 2**). When inspecting the structures of a single subunit of each multimerization domain-containing design alone, each design appeared to have the multimerization domain portion accessible, suggesting multimerization should be possible. Predictions using the multimer parameter supported the ability of the structures to form multimers of 3-mers, 7-mers, and 24-mers for mSb-Foldon, mSb-IMX313, and mSb-Ferritin, respectively, as expected. Additionally, these structures also show the varying geometry of each multimerization domain: foldon generates a “bouquet”-like geometry, IMX313 generates a circular ring geometry, and ferritin generates a spherical geometry. Finally, these data suggest approximate protein particle sizes of 9.5 nm W/L x 6.6 nm H for mSb-Foldon, 11.5 nm W/L x 7.5 nm H for mSb-IMX313, and 22 nm W/L and H for mSb-Ferritin. Therefore, these predicted multimer sizes support APC-independent lymph node drainage (10-100 nm [11–13]) as a potential mechanism towards increasing immunogenicity for IMX313 and ferritin multimers.

**Figure 2.**
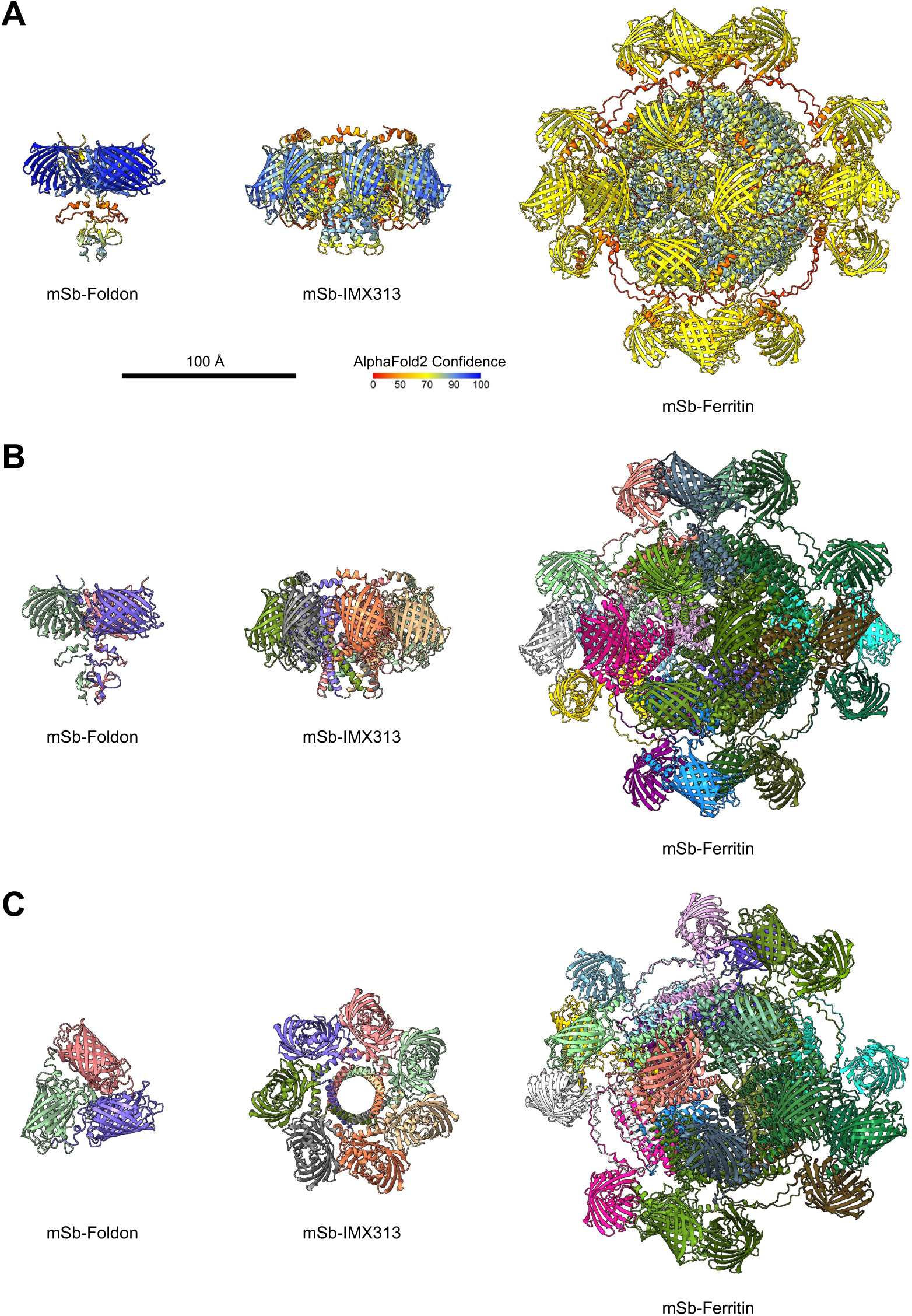
Structural prediction of multimeric mSb designs. mSb multimers were predicted after C-terminal addition of multimerization domains and removal of the IgK signal peptide using AlphaFold2 (mSb-Foldon, left; mSb-IMX313, middle; mSb-Ferritin, right). A) AlphaFold2 confidence of the predicted structures (side-view). B/C) Structures with each monomer individually coloured (side-view, B; top-view, C). Scale bar applies to all structures.

### In vitro evaluation of antigen designs

Secreted monomeric and secreted multimeric designs were then evaluated *in vitro* for expression/secretion and expression/secretion/multimerization, respectively (**Figure 3**). For these experiments, mSb-Strep (intracellular) served as a negative control for evaluating secretion. K562 cells were transfected with mSb-Strep (intracellular), Igκ-mSb-Strep (secreted monomeric) mRNA LNP or mock transfected (negative control for transfection/mSb expression) with or without addition of protein transport inhibitor cocktail (PTIC), and then mSb fluorescence was evaluated by flow cytometry (**Figure 3A**). PTIC contains brefeldin A and monensin which are well-established inhibitors of the export pathway in mammalian cells [49,50]. Addition of PTIC had no effect on mock transfected negative control cells or mSb-Strep mRNA LNP transfected cells. Conversely, addition of PTIC in Igκ-mSb-Strep mRNA LNP transfected cells resulted in increased mSb fluorescence. Given that inhibition of secretion led to intracellular accumulation of mSb, these data indicate the inclusion of the N-terminal Igκ signal peptide successfully leads to secretion in our designs. All mRNA LNP successfully led to mSb expression - transfection resulted in mSb fluorescent positivity of 99.0% for mSb-Strep, 85.7% for Igκ-mSb-Strep, 98.5% for Igκ-mSb-Foldon, 99.0% for Igκ-mSb-IMX313, and 67.8% for Igκ-mSb-Ferritin, compared to 0.1% positivity in mock transfected negative controls (**Figure 3B**).

**Figure 3.**
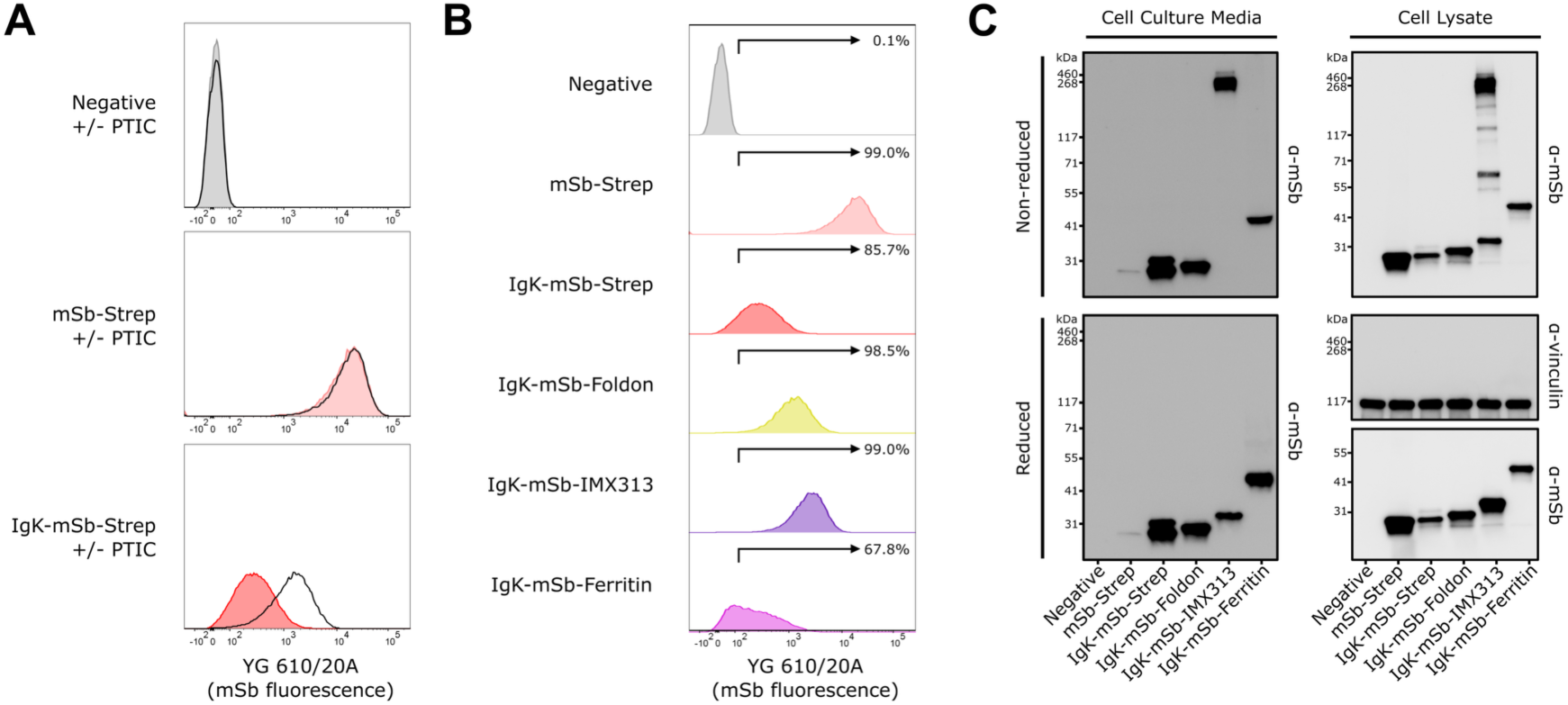
*In vitro* validation of secreted monomeric and secreted multimeric mSb-encoding mRNA LNP vaccines. A) Flow cytometry analysis of K562 cells transfected with mRNA LNP with and without addition of protein transport inhibitor cocktail (PTIC). K562 cells were transfected with 1.25 μg of mRNA LNP. At 16 hours following transfection, PTIC was added to respective conditions (black line histograms). At 24 hours following transfection (i.e. 8 hours following addition of PTIC), cells were collected and evaluated for mSb fluorescence by flow cytometry. Events were gated to isolate live singlet cells. Data is representative of 2 replicates (1 replicate shown). See Supplemental Figure S1 for gating strategy. B) Flow cytometry analysis following transfection of K562 cells with mRNA LNP. K562 cells were transfected with 1.25 μg of mRNA LNP. At 24 hours following transfection, cells were collected and evaluated for mSb fluorescence by flow cytometry. Events were gated to isolate live singlet cells. Data is representative of 2 replicates (1 replicate shown). See Supplemental Figure S1 for gating strategy. C) Western blot analysis of K562 cells transfected with 1.25 μg mRNA LNP. At 24 hours following transfection, cells (for preparing cell lysate) and culture media were collected. Samples were run on SDS-PAGE in reducing or non-reducing conditions, transferred to nitrocellulose, probed via anti-mSb and anti-vinculin (loading control) antibodies. Blots were then probed with appropriate HRP-conjugated secondary antibodies, then developed and imaged.

mSb expression, secretion, and multimerization were evaluated by western blot analysis of cell lysate and cell culture media samples (**Figure 3C**). Cell lysate samples run under reducing conditions showed the expected banding of all mSb designs (Igκ = 2.4 kDa, mSb = 26.6 kDa, StrepTagII = 1.1 kDa, foldon = 3.1 kDa, IMX313 = 6.2 kDa, ferritin = 20.0 kDa). Cell culture media samples run under reducing conditions similarly demonstrated expected banding of all mSb designs. Of note, cell culture media of Igκ-mSb-Strep appeared as 2 similarly sized bands, possibly reflecting a species with post-translational modifications associated with the secretion pathway. Cell culture media of mSb-Strep had only a faintly present band, likely representing release of protein from cells which died/lysed in the period between transfection and collection. Thus, these data further support that the N-terminal Igκ signal peptide successfully leads to antigen secretion, as observed by flow cytometry, and demonstrates successful secretion of Igκ-mSb-Foldon, Igκ-mSb-IMX313, and Igκ-mSb-Ferritin. Given that foldon, IMX313, and ferritin-based multimerization results in 3-mer, 7-mer, and 24-mer products, respectively, a corresponding 3x, 7x, and 24x shift in molecular weight is expected. Comparing reducing and non-reducing conditions, a shift in band size was observed for Igκ-mSb-IMX313, corresponding to formation of a 7-mer product (32 kDa * 7 = 224 kDa). Additionally, the presence of only the 7-mer band in the cell media sample of Igκ-mSb-IMX313 suggests complete multimerization of these monomers prior to secretion, and that the multimer is relatively stable in the extracellular environment. The presence of multiple other bands in the cell lysate sample of Igκ-mSb-IMX313 possibly reflects a stepwise nature of multimer formation. No bands corresponding to multimers were observed for Igκ-mSb-Foldon or Igκ-mSb-Ferritin. Under non-reducing or reducing conditions, multimeric and monomeric forms, respectively, have been observed for foldon-based [51], IMX313-based [22,52], and ferritin-based multimers [51]. Therefore, under conditions permissive to multimer detection, we find no evidence of multimerization for IgK-mSb-Foldon or IgK-mSb-Ferritin.

Next, surface-displayed designs were evaluated *in vitro* for expression and surface-display (**Figure 4**). For these experiments, mSb-Strep (intracellular) served as a negative control for evaluating surface-display. HEK293T cells were transfected with mSb-Strep, Igκ-mSb-TM(PDGFR)-Strep, or Igκ-mSb-TM(B7)-Strep pDNA or mock transfected (negative control for transfection/mSb expression) and then mSb fluorescence and mSb surface staining was evaluated by flow cytometry (**Figure 4A**). Transfection resulted in mSb fluorescent positivity of 69.6% for mSb-Strep, 81.1% for Igκ-mSb-TM(PDGFR)-Strep, and 70.7% for Igκ-mSb-TM(B7)-Strep, compared to 0.1% positivity in mock transfected negative controls, indicating all designs successfully lead to mSb expression. Alternatively, surface-staining was detected at a positivity of 0.3% for mSb-Strep, 98.2% for Igκ-mSb-TM(PDGFR)-Strep, and 98.1% for Igκ-mSb-TM(B7)-Strep, compared to 0.3% positivity in mock transfected negative controls. These data showed that inclusion of the N-terminal Igκ signal peptide with a PDGFR or B7 TM domain results in successful surface-display.

**Figure 4.**
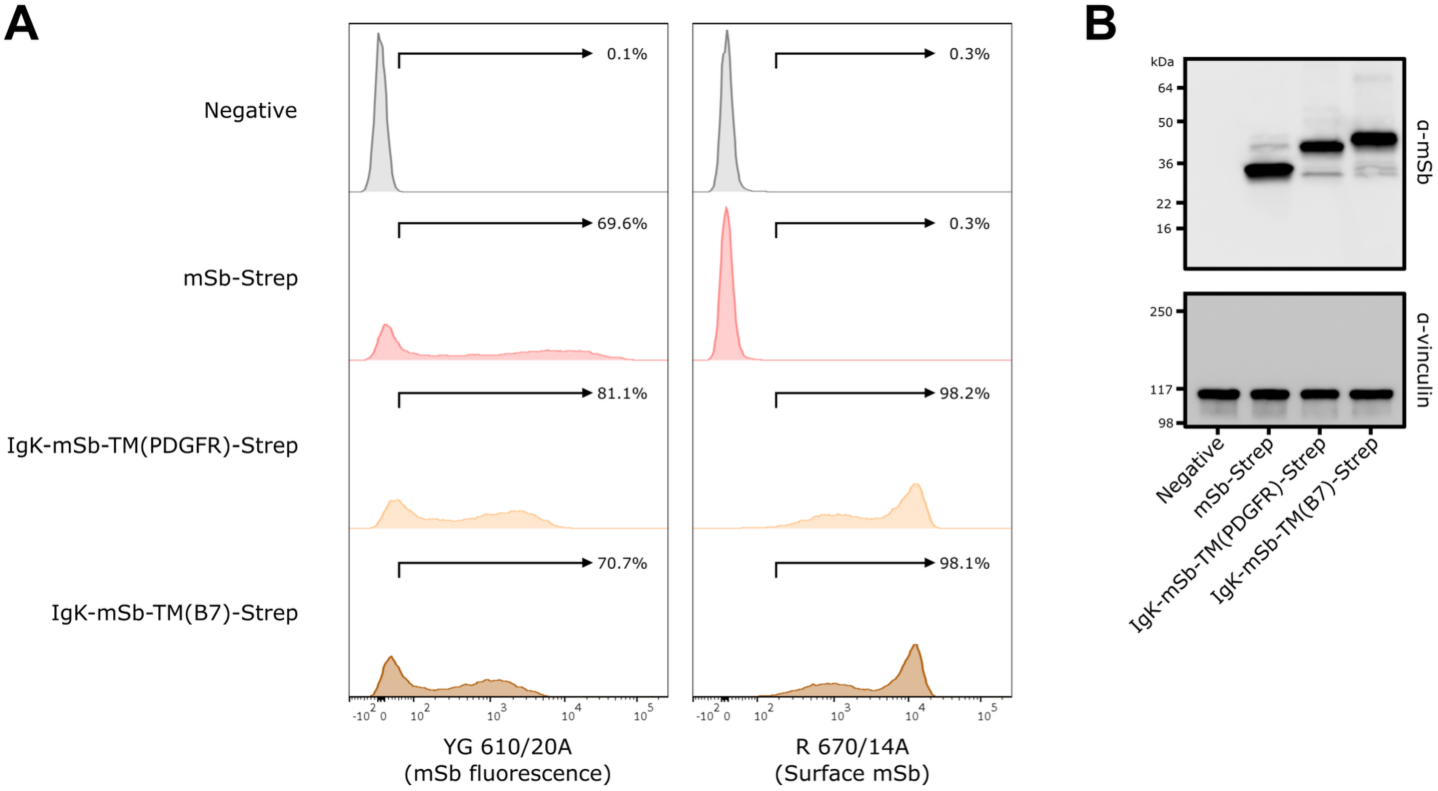
*In vitro* validation of surface-displayed mSb-encoding pDNA constructs used to generate mRNA LNP vaccines. A) Flow cytometry analysis of HEK293T/17 cells transfected with 2.5 μg of pDNA. At 24 hours following transfection, cells were collected and stained with anti-mSb and then AlexaFluor647-conjugated secondary antibodies and then evaluated for mSb fluorescence and surface staining by flow cytometry. Events were gated to isolate live singlet cells. Data is representative of 2 replicates (1 replicate shown). See Supplemental Figure S2 for gating strategy. B) Western blot analysis of HEK293T/17 cells transfected with 2.5 μg pDNA. At 24 hrs post-transfection, cells (for preparing cell lysate) were collected. Cell lysates were prepared and run on SDS-PAGE in reducing conditions, transferred to nitrocellulose, and then probed via anti-mSb and anti-vinculin (loading control) antibodies. Blots were then probed with appropriate HRP-conjugated secondary antibodies, then developed and imaged.

As a secondary validation of expression, western blot analysis was performed on cell lysates of mSb-Strep, Igκ-mSb-TM(PDGFR)-Strep or Igκ-mSb-TM(B7)-Strep transfectants (**Figure 4B**). A dominant band corresponding to expected sizes for all mSb designs was observed (Igκ = 2.4 kDa, mSb = 26.6 kDa, TM(PDGFR) = 5.4 kDa, TM(B7) = 7.7 kDa, StrepTagII = 1.1 kDa). Igκ-mSb-TM(B7)-Strep was selected for further evaluation given the ∼7.6-fold longer half-life of B7-based surface-display compared to PDGFR-based surface-display [53].

### In vivo evaluation of antigen designs

Next, to determine the relative immunogenicity of the various mSb-encoding designs, an *in vivo* prime-boost vaccination study was performed in C57BL/6-A0201 mice, comparing mRNA LNP vaccines of the following designs: Igκ-mSb-Strep, Igκ-mSb-TM(B7)-Strep, Igκ-mSb-Foldon, Igκ-mSb-IMX313, or Igκ-mSb-Ferritin (**Figure 5A**). Of note, the mice used were transgenic for HLA-A*02:01^+^ so as to allow future assessment of MHC class I epitopes presented by this allele, which is beyond the scope of the present study. At experimental endpoint (14-days following boost vaccination), blood samples were collected, and serum was isolated. To assess immunogenicity following vaccination, an indirect ELISA using recombinant mSb-coated plates was performed for serum of each mouse (**Figure 5B**). All vaccines elicited responses compared to negative control serum samples derived from the same mouse strain (p<0.05; ANOVA), except Igκ-mSb-Ferritin. No significant difference was observed between Igκ-mSb-Strep and Igκ-mSb-TM(B7)-Strep (p≥0.05), indicating secretion and surface-display are equally immunogenic in this context. Similarly, Igκ-mSb-Foldon or Igκ-mSb-Ferritin showed no difference (p≥0.05) in antigen-specific IgG relative to the Igκ-mSb-Strep. Assuming this lack of increased antigen-specific IgG was due to lack of multimerization, these findings are agreement with our *in vitro* observations. Alternatively, Igκ-mSb-IMX313 generated ∼5-fold greater antigen specific IgG compared to its monomeric counterpart (p<0.05). These data indicate that high rates of antigen self-multimerization can be achieved from mRNA LNP, translating to improved antibody responses, and suggest that IMX313 may be well-suited as a multimerization domain in the context of mRNA LNP vaccination.

**Figure 5.**
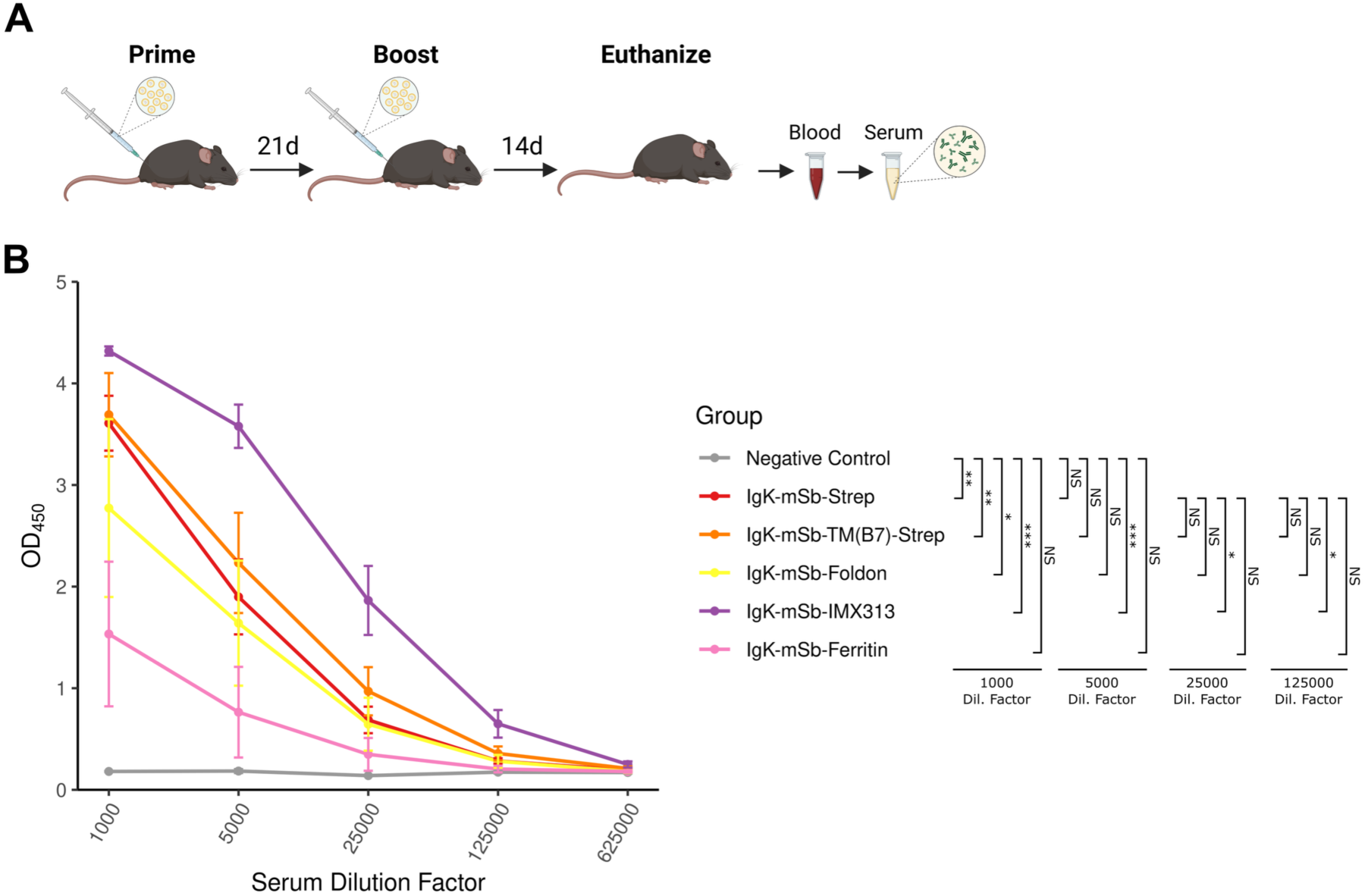
*In vivo* evaluation of secreted monomeric, secreted multimeric, and surface-displayed mSb-encoding mRNA LNP vaccines. A) *In vivo* study design for testing of mSb-encoding mRNA LNP vaccination in C57BL/6J-A0201 mice (*n* = 4 mice/group). Figure was made using BioRender.com. B) Indirect ELISA analysis of negative control C57BL/6J-A0201 serum (*n* = 3) and endpoint serum isolated from vaccinated mice of each mRNA LNP vaccine type (*n* = 4 mice/group). Immulon 2HB plates were coated overnight with 100 μl/well of 2 μg/ml recombinant mSb, blocked, then individual mouse serum was added at various dilutions and incubated overnight, then plates were incubated with HRP-conjugated donkey anti-mouse IgG secondary antibodies, developed using TMB, and the OD_450_ was read. Points represent the mean, and error bars represent SEM. Significance was determined using a one-way ANOVA with Tukey’s HSD.

## Discussion

Compared to protein vaccines, reports of the use of multimerization domains in mRNA LNP vaccines are limited. BNT162b1 was a SARS-CoV-2 mRNA LNP vaccine candidate developed by Pfizer/BioNTech, which encoded the Spike protein receptor binding domain (RBD) with a foldon multimerization domain to generate trimeric multimers for secretion and was evaluated in a phase I/II clinical trial (though ultimately not rolled out due to increased reactogenicity compared to BNT162b2, despite similar level of antibody responses) [54,55]. While this candidate demonstrated multimerization *in vitro* following mRNA LNP transfection [18], there was no *in vivo* evaluation compared to a secreted monomeric control that would enable conclusions as to the possible benefits of multimer formation. However, similar secreted trimeric foldon-based RBD designs delivered as mRNA LNP have been directly compared *in vivo* to monomeric equivalents (secreted monomeric RBD, as well as surface-displayed native spike) and demonstrated superior neutralizing titer and strain cross-reactivity in mice [56]. Similarly, ferritin-based multimerization of trimeric RBD subunits significantly increased neutralization titers and IFN-γ reactive cells compared to trimeric RBD subunits without the ferritin domain in mRNA LNP vaccinated mice [51]. Alternatively, others report only limited proportions of properly folded multimeric antigen (0.7%) for ferritin-based designs delivered as mRNA, as well as reduced *in vitro* and *in vivo* expression and reduced immunogenicity of such designs compared to surface-displayed comparators [57]. In our data, foldon- and ferritin-based designs did not multimerize to a detectable level and did not improve immunogenicity. Thus, the mechanisms allowing foldon and ferritin domains to multimerize and enhance immunogenicity of mRNA LNP vaccines in some contexts and not others require further investigation.

To the best of our knowledge, our data represents the first evaluation of the IMX313 multimerization domain in the context of mRNA LNP vaccination. Of particular note, in this study, the IMX313-containing design, Igκ-mSb-IMX313, was found to multimerize *in vitro* and increase antibody responses *in vivo*, while foldon- and ferritin-containing designs (Igκ-mSb-Foldon and Igκ-mSb-Ferritin) did not. These data suggest that, at least for some antigens, the IMX313 multimerization domain may be superior to foldon and ferritin in the setting of mRNA LNP vaccines. We propose three possible explanations for this finding: higher abundance level, greater inter-monomer affinity, and lack of structural interference. Firstly, the IMX313-containing design had greater abundance following transfection than any other secreted design. It is possible this enhanced abundance level directly drove, at least in part, the observed enhanced immunogenicity. Alternatively, effective multimerization may be dependent on a critical threshold of abundance. Increased intracellular antigen concentration would promote the likelihood of interactions between subunits, necessary for multimer formation. It is unclear as to why the IMX313-containing design had increased abundance levels, but this may be related to increased solubility, as has been reported for other protein fusions such as MBP fusions [58]. Further, high abundance levels may be particularly important for secreted antigen, given the limited time between antigen translation and secretion (after which antigen concentration will much lower in the extracellular environment and therefore less likely to multimerize). Alternatively, the IMX313 domain may have greater inter-monomer affinity, and therefore form more stable multimers that remain bound together during and following secretion. Stable multimer formation may also be more important in situations of lower cellular antigen abundance that would reduce the likelihood of reformation following dissociation. Lastly, there may be structural interference for specific combinations of antigen of interest and multimerization domain (i.e. mSb in combination with IMX313 may have allowed uninhibited multimer formation, whereas there may have been occlusion of the foldon and ferritin multimer domain by the mSb portion of the chimeric protein). While we performed structural predictions with the goal of selecting candidate mSb/multimer domain combinations without evidence of such issues, the imperfect nature of these predictions makes this remain a possible consideration.

Future work may expand upon our results. Firstly, we selected mSb as a representative antigen candidate of interest, given its fluorescence makes it readily detected, however, as mentioned above, alternative antigens may be better suited for some multimerization domains, therefore these findings should be evaluated in additional contexts. Similarly, we selected 3 widely used multimerization domains (foldon, IMX313, and ferritin) with varying geometries, sizes, and orders of multimerization (3-, 7-, 24-mers). Future studies evaluating additional multimerization domains in the context of mRNA LNPs may reveal commonalities between those best-suited for mRNA LNP vaccine application. In addition, increased immunogenicity from multimerization does not necessarily directly correlate with increased protection. The capacity of multimeric designs to target key regions of pathogen-derived antigens and similarly evoke enhanced neutralization and enhanced protection should be explored in future work. Lastly, we used secreted multimeric antigen designs and focused on antibody responses induced by these vaccines, given that antibody titer/neutralizing titer are the primary correlates of protection for vaccines against many infectious diseases [59] and antibody responses are most suitable for our applications of interest. However, a limitation of our study is that vaccine-induced T cell responses were not evaluated, and can be critical for protection in some disease contexts, such as some intracellular pathogens and cancer. Assessment of the impact of multimerization on such responses may be beneficial for certain vaccine applications. In principle, surface-displayed antigen may generate better CD8^+^ T cell responses as the antigen is retained by the cell and can be processed and presented on MHC I. Therefore, developing multimeric forms of surface-displayed antigen may have significant potential, inducing stronger antibody responses through possible synergistic effects of surface-display and multimerization, while also being retained for MHC I presentation to induce CD8^+^ T cell responses, and should be explored in future studies.

## Conclusion

In conclusion, our data supports the applicability of multimerization domains in mRNA LNP vaccine design, and highlights IMX313 as a potential preferential domain for mRNA LNPs. Further research exploring factors that lead to successful multimerization and enhancement of immunogenicity in the mRNA LNP setting would be informative for future vaccine development efforts.

## Materials & Methods

### Multimer structural predictions

Alphafold 2.3.1 [48] was used to predict structures for multimerized mSb constructs with foldon (3-mer), IMX313 (7-mer), and ferritin (24-mer). Each prediction run used one NVIDIA RTX A6000 GPU, 32 CPUs, and 220GB RAM. The “-m multimer” parameter was used. For foldon and IMX313 versions, protein reference files were provided with 3 and 7 copies of the construct sequence, respectively. For ferritin, to make the prediction computationally tractable, 24 copies of the ferritin sequence were predicted in a multimer run, and separately, 1 copy of the mSb-Ferritin construct was predicted. The highest confidence model for each prediction run was selected. For mSb-Ferritin, the alpha-carbons of the ferritin portion of the monomer were aligned to the structure of each of the multimer 24mer ferritins using ChimeraX, generating the final multimerized structure.

### Plasmid DNA generation

DNA sequences for all plasmids used in this study are listed in **Supplementary Data File 1**. For design of golden gate assembly parts, coding sequences were codon-optimized using GenSmart (GenScript) and then manually edited to remove uridine residues in the wobble position of two-family codons. RNA structures were predicted by RNAFold [60] and coding sequences were modified as needed to maintain an optimal RNA structure in the 5’ UTR. All golden gate assembly parts were domesticated to remove Type IIs restriction sites and then incorporated into pNL and/or pLZ1/4. To create the pRNA series of plasmids, pRNA-Destination plasmids was combined with appropriate pLZ plasmids and assembled in a *PaqC*I-based golden gate assembly reaction performed using manufacturers protocols. In brief, reactions contained 50 ng of destination vector and equimolar amounts of pLZ, 10 U *PaqC*I, 400 U of T4 DNA ligase, and 5 pmoles of *PaqC*I activator in T4 DNA ligase buffer and were cycled 30 times for 37°C for 5 mins then 16°C for 5 mins. All pDNA constructs were sequence verified. All enzymes were purchased from New England Biolabs. All golden gate assembly parts were ordered from Integrated DNA Technologies, GenScript, or Twist Bioscience.

NEB stable *E. coli* (New England Biolabs, Cat. #C3040H) were used for propagation of DNA. *E. coli* was transformed following manufacturer protocols. *E. coli* was grown in LB broth at 30°C, shaking at 250 rpm. For solid medium, LB broth was supplemented with Bacto agar (1.5% [w/v]) (BD, Cat. #214010) and further supplemented with 100 μg/ml carbenicillin or 50 μg/ml kanamycin, as appropriate. For blue/white screening, media was further supplemented with 1 mM IPTG and 0.1 mg/ml of X-Gal (prepared as a 20 mg/ml stock solution in DMF). All pDNA was quantified by Qubit assay and stored at -20°C.

### *mRNA generation via in vitro transcription* (IVT)

pDNA template was linearized in preparation for IVT by restriction digest using BspQI (NEB, Cat. #R0712L) in 1x NEBuffer 3.1 (NEB, Cat. #B7203) with incubation at 50°C for ≥1 hr. Linearized pDNA was then column-cleaned using the Monarch PCR & DNA cleanup kit (NEB, Cat. #T1030) following manufacturer’s protocols and eluting with 50°C elution buffer. IVT was performed by combining 1000 ng of pDNA with RNA pol reaction buffer (NEB, Cat. #B9012) (1x final), magnesium acetate (Sigma, Cat. #63052-100ML) (12.5 mM final), Triton X-100 (Sigma, Cat. #93443) (0.002% final), ATP (NEB, Cat. #N0450S) (5 mM final), GTP (NEB, Cat. #N0450S) (5mM final), CTP (NEB, Cat. #N0450S) (5 mM final), N1-methyl-pseudouridine-TP (TriLink Biotechnologies, Cat. #N-1081-1) (5mM final), CleanCap Reagent AG (TriLink Biotechnologies, Cat. #N-7113-5) (4 mM final), RNase inhibitor (NEB, Cat. #M0314L) (10 U/μl final), DTT (Thermo Scientific, Cat. #707265ML) (5 mM final), pyrophosphatase (NEB, Cat. #M0361L) (0.0025 U/μl final), and T7 polymerase (NEB, Cat. #M0251) (5 U/μl final) in a total reaction volume of 20 μl (brought to 20 μl with UltraPure dH_2_O [Invitrogen, Cat. #10977023]) and incubating at 37°C for 4-6 hrs. After incubation, 28 μl of UltraPure dH_2_O and 2 μl of DNAse I (NEB, Cat. #M0303S) was added then samples were incubated at 37°C for 15 mins. Resulting mRNA was then column-cleaned using the Monarch RNA cleanup kit (NEB, Cat. #T2040) following manufacturer’s protocols with 2 additional wash steps and 5 min RT incubation before centrifugation steps after addition of washing or elution buffers. After clean-up, mRNA was quantified using a Qubit BR RNA assay kit (Thermo Scientific, Cat. #Q10211) and integrity was verified by gel electrophoresis. mRNA was stored at -80°C.

### mRNA LNP encapsulation and quantification

For encapsulation, mRNA was diluted to 1 mg/ml with UltraPure dH_2_O (Invitrogen, Cat. #10977023) in a total volume of 22 μl. mRNA was then encapsulated using the NanoAssemblr Spark Hepato9 siRNA kit (Cytiva, Cat. #NWS0009) following manufacturer’s protocols and using 16 μl of Spark Nanoparticle mix. Resulting mRNA LNP was evaluated for concentration and encapsulation efficiency using the Quant-IT RiboGreen RNA assay kit (Thermo Scientific, Cat. #R11490) based on manufacturer’s protocols with or without the addition of Triton-X 100 (Sigma, Cat. #93443) (2% final) and measured on a VICTOR2 plate reader. All mRNA LNPs were determined to have encapsulation efficiencies of ≥93.6%. Concentration of encapsulated mRNA was used for all further assays. mRNA LNP of each set were generated within 1 day of each other and stored at 4°C for ≤7 days before use.

### Cell culture

HEK293T/17 cells were cultured in cDMEM (complete DMEM; DMEM [Gibco, Cat. #11995073] supplemented with 10% heat-inactivated FBS (Gibco, Cat. #A3840302 or Cat. #12484028), 1X GlutaMAX [Gibco, Cat. #35050061], 1X Penicillin-Streptomycin [Gibco, Cat. #15140122], 1X MycoZap [Lonza, Cat. #VZA-2031]). HEK293T/17 cells were dissociated using TrypLE Express (Gibco, Cat. #12604013) for routine subculturing or using TrypLE Express or enzyme-free cell dissociation buffer (Gibco, Cat. #13151014) following transfection (enzyme-free cell dissociation buffer was used to dissociate cells following transfection for experiments requiring surface proteins to be preserved). K562 cells were cultured in cRPMI (complete RPMI; RPMI 1640 [Gibco, Cat. #11875119] supplemented with 10% heat-inactivated FBS, 1X GlutaMAX, 1X Penicillin-Streptomycin, 1X MycoZap, 10 mM HEPES [Gibco, Cat. #15630080], 1 mM sodium pyruvate [Gibco, Cat. #11360070], and 55 μM 2-mercaptoethanol [Gibco, Cat. #21985023]). All cells were grown in an incubator at 37°C, 5% CO_2_. Cells were cryopreserved in 90% FBS/10% DMSO in vapor-phase liquid nitrogen. HEK293T/17 and K562 cells were purchased from ATCC.

### Cell transfection with pDNA and mRNA LNP

For mRNA LNP transfections of K562 cells, cells were seeded at a density of 600,000 cells/ml in 6-well tissue culture-treated plates (2.5ml/well) in cRPMI supplemented with 1 μg/ml ApoE4 (Peprotech, Cat. #350-04). mRNA LNP were adjusted to 8.33 ng/μl with D-PBS (MgCl^-^, CaCl^-^) (Gibco, Cat. #14190250) and then cells were transfected by addition of 1.25 μg of mRNA LNP per well (150 μl/well) dropwise, then plates were swirled and rocked to disperse complexes. For mock transfected negative controls, D-PBS (MgCl^-^, CaCl^-^) was used in place of mRNA LNP. For conditions with protein transport inhibition, protein transport inhibitor cocktail (Invitrogen, Cat. #00-4980-83) was diluted in media and added to transfectants at ∼1x final concentration at 16 hours following transfection.

For pDNA transfections, HEK293T/17 cells were seeded at a density of 140,000 cells/cm^2^ in 6-well tissue culture-treated plates (540,000 cells/ml, 2.5 ml/well) (Corning, Cat. #353046). Transfections were performed at ∼16.5 hours following seeding. Based on manufacturer’s recommendations, pDNA and buffer EB (Qiagen, Cat. #19086) (to normalize total pDNA and buffer EB volume depending on pDNA concentration) were combined with TransIT-LT reagent (Mirus Bio, Cat. #MIR2305) at a ratio of 3 μl of TransIT-LT per 1 μg of pDNA per 108 μl of total volume in OptiMEM (Gibco, Cat. #3195062) and incubated at RT for ∼20 mins to allow complex formation. For mock transfected negative controls, buffer EB alone was used in place of buffer EB/pDNA. Following complex formation, a volume corresponding to 2.5 μg of pDNA was added dropwise per well, and plates were swirled and/or rocked to disperse complexes.

### Flow cytometry

For assessment of mSb fluorescence alone in transfectants, at ∼24 hours following transfection, cells were transferred to microfuge tubes, centrifuged at 300 x *g* (4°C), supernatant was removed, and cells were resuspended in D-PBS (MgCl^-^, CaCl^-^) (Gibco, Cat. #14190250) and counted on an EVE automated cell counter. A volume equivalent to 1,000,000-1,500,000 cells was then centrifuged at 300 x *g* (4°C), supernatant was removed, and cells were resuspended in 500 μl FACS buffer (5% FBS in D-PBS). Cells were centrifuged at 300 x *g* (4°C), supernatant was removed and then cells were resuspended in 400 μl of FACS buffer with 0.1 μg/ml DAPI (Sigma, Cat. #D9542). Cells were strained through a 0.35 μm FACS tube cap (Corning, Cat. #352235), placed on ice, and analyzed on a BD LSRFortessa flow cytometer.

For assessment of mSb fluorescence and surface mSb expression in transfectants, at ∼24 hours following transfection, cells were dissociated using 1 ml/plate of enzyme-free cell dissociation buffer (Gibco, Cat. #13151014) (to preserve surface proteins). Cells were transferred to microfuge tubes, centrifuged at 300 x *g* (4°C), supernatant was removed, and cells were resuspended in D-PBS (MgCl^-^, CaCl^-^) (Gibco, Cat. #14190250) and counted on an EVE automated cell counter. A volume equivalent to ∼500,000 cells was then centrifuged at 300 x *g* (4°C), supernatant was removed, and then cells were stained with 100 μl of 1/1000 anti-mSb monoclonal mouse IgG antibody (OriGene, Cat. #TA180039) (diluted 1/1000 in FACS buffer [5% FBS in D-PBS]) for 30 mins at 4°C. Cells were then centrifuged at 300 x *g* (4°C), supernatant was removed, and cells were resuspended in 500 μl FACS buffer twice. Cells were then centrifuged at 300 x *g* (4°C), supernatant was removed, and cells were then stained with 100 μl of 1/200 diluted AlexaFluor647-conjugated goat anti-mouse IgG polyclonal antibody (Biolegend, Cat. #405322) (diluted 1/200 in FACS buffer) for 30 mins at 4°C. Cells were then centrifuged at 300 x *g* (4°C), supernatant was removed, and cells were resuspended in 500 μl FACS buffer twice. Cells were then centrifuged at 300 x *g* (4°C), supernatant was removed, and cells were resuspended in 400 μl of FACS buffer with 0.1 μg/ml DAPI (Sigma, Cat. #D9542). Cells were strained through a 0.35 μm FACS tube cap (Corning, Cat. #352235), placed on ice, and analyzed on a BD LSRFortessa flow cytometer.

### Western blot

For assessment of transfectant media in western blot, at ∼24 hours following transfection, media from K562 cells (containing cells, as K562 cells are suspension cells) was collected from wells by transferring to a microfuge tube. Samples were centrifuged at 300 x *g* for 5 mins to pellet cells and supernatant was transferred to a new tube twice. cOmplete EDTA-free protease inhibitor (Roche, Cat. 11873580001) was then added to 1x final. Media samples were then stored at 4°C (short-term, i.e. 1 day) for multimer experiments or - 20°C for all other experiments until western blot analysis.

For assessment of transfectant cell lysate in western blot, cells were collected at ∼24 hours following transfection. For HEK293T/17 cells, cells were dissociated using enzyme-free cell dissociation buffer (Gibco, Cat. #13151014) (to preserve surface proteins). Cells were then transferred to microfuge tubes, centrifuged at 300 x *g*, supernatant was removed, and cells were resuspended in D-PBS (MgCl^-^, CaCl^-^) (Gibco, Cat. #14190250) and counted on an EVE automated cell counter. For K562 cells, cells (in media, as K562 cells are suspension cells) were transferred to microfuge tubes, centrifuged at 300 x *g*, supernatant was removed, and cells were resuspended in D-PBS (MgCl^-^, CaCl^-^) (Gibco, Cat. #14190250) and counted on an EVE automated cell counter. A volume equivalent to ∼500,000 cells was then centrifuged at 300 x *g*, supernatant was removed, and cells pellets were stored at -20°C. Cell pellets were then thawed and resuspended in 100 μl of RIPA buffer (Thermo Scientific, Cat. #89900) with cOmplete EDTA-free protease inhibitor (Roche, Cat. #11873580001) added to 1-1.25x final. Samples were then incubated on ice for 15 mins to lyse, then centrifuged at 14,000 x *g* for 15 mins at 4°C. Supernatant (corresponding to prepared cell lysate) was then transferred to a new tube. Lysed cell pellets and prepared cell lysates were stored at 20°C. As needed, lysed cells pellets were re-centrifuged, and supernatant was re-transferred to generate more prepared cell lysate sample.

Transfectant media or prepared cell lysate samples were then combined with a 1:3 volume ratio with Laemmli buffer (4x) (BioRad, Cat. #1610747) supplemented with (reducing) or without (non-reducing) 2-mercaptoethanol (1.42 M final) (BioRad, Cat. #1610710), or with (reducing) DTT (50 mM final). Samples were heated (85°C for sample sets with multimers, 95°C for 5 mins for same sets without multimers) and then 5 μl were loaded per lane onto a 4-15% mini-PROTEAN TGX polyacrylamide gel (BioRad, Cat. #4561085 or 4561086) and ran at 50V for 15-30 mins, then 150V for 35-60 mins in 1x Tris/Glycine/SDS running buffer (BioRad, Cat. #1610732). Proteins were then transferred to nitrocellulose blots in transfer buffer (1x Tris/Glycine [BioRad, Cat. #1610734], with 20% methanol) at 0.30 A for 1.33-1.5 hrs on ice. After transfer, blots were cut as needed, washed 1x in 10 ml TBS-T (TBS with 0.1% Tween-20) then blocked with 10 ml of EveryBlot blocking buffer (BioRad, Cat. #12010020) for 10 mins at RT. Blots were then washed 2x in 10 ml TBS-T and then probed with 10 ml of mouse anti-StrepTagII monoclonal antibody (diluted to 1/5000 in EveryBlot blocking buffer) (LSBio, Cat. #LS-C413454), mouse anti-mSb monoclonal antibody (diluted 1/2000 in EveryBlot blocking buffer) (OriGene, Cat. #TA180039), or rabbit anti-vinculin monoclonal antibody (diluted 1/5000 in EveryBlot blocking buffer) (Abcam, Cat. #ab129002) overnight at 4°C. Blots were then washed 4x in 10 ml TBS-T and then probed with 10 ml of HRP-conjugated goat anti-mouse IgG polyclonal antibody (diluted 1/10,000 in EveryBlot blocking buffer) (Jackson Immunology, Cat. #115035003) or HRP-conjugated goat anti-rabbit IgG polyclonal antibody (diluted 1/10,000 in EveryBlot blocking buffer) (Jackson Immunology, Cat. #111035003), as appropriate, for 1 hr at RT. Blots were then washed 4x in 10 ml TBS-T, developed using clarity western ECL substrate (BioRad, Cat. #1705060) and then imaged on a BioRad ChemiDoc imaging system. SeeBluePlus2 pre-stained protein standard (Invitrogen, Cat. #LC5925) or HiMark pre-stained protein standard (Invitrogen, Cat. #LC5699) was ran alongside samples to evaluate molecular weight.

### Mice

Female C57BL/6J-A0201 (C57BL/6J-Mcph1^Tg(HLA-A2.1)1Enge^/J [Jackson strain #003475]) mice were purchased from Jackson Laboratory and allowed to acclimate for ≥7 days following arrival. Mice were 7-8 weeks of age at the beginning of experiments. All experiments involving animals were performed within the BC Cancer Animal Resource Centre (ARC) under specific pathogen-free conditions. All experiments involving animals were performed with approval from the UBC Animal Care Committee (A21-0198, approved Oct 26, 2021).

### mRNA LNP vaccinations

mRNA LNP vaccines were adjusted to 50 ng/μl using D-PBS (MgCl^-^, CaCl^-^) (Gibco, Cat. #14190250). mRNA LNP vaccines were then placed on ice, transferred to the ARC, and kept on ice/at 4°C until injection. Immediately prior to injection, mRNA LNP vaccines were warmed to RT. Each mouse was injected intramuscularly with 20 μl of mRNA LNP vaccine (1 μg/mouse). All vaccinations were performed as a prime-boost regimen, with boost vaccinations 21 days following prime vaccinations and with experiment endpoint 14 days following boost vaccinations.

### Serum isolation

For endpoint plasma/serum, immediately following euthanasia at experimental endpoint by isoflurane followed by CO_2_ inhalation, whole blood was collected via cardiac puncture with a 25G needle. Whole blood was placed in an SST microtainer tube (BD, Cat. #365967), clotted for 30-60 mins at RT, and then placed on ice/at 4°C. To isolate serum, whole blood was then centrifuged at 2000 x *g* for 10 mins at 4°C and the supernatant/upper phase (corresponding to serum) was then transferred to a new tube and stored at -80°C. For testing of pooled serum samples, serum was pooled by combining equal volumes of serum from each mouse within each experimental group.

### Enzyme-linked immunosorbent assay (ELISA)

Immulon 2HB (Thermo Scientific, Cat. #14-245-61) were coated by addition of 100 μl/well of 2 μg/ml of recombinant mStrawberry (OriGene, Cat. #TP790044) in ELISA carbonate coating buffer (Invitrogen, Cat #CB01100) and incubation overnight at 4°C. Plates were then washed 3 times with 200 μl/well of 1x wash buffer (Invitrogen, Cat. #WB01). Wells were then blocked by addition of 200 μl/well of 1x ELISA assay buffer (Invitrogen, Cat. #DS98200) and incubation at RT for 3 hrs. Plates were then washed 3 times with 200 μl/well of 1x wash buffer. Next, 100 μl/well of individual serum samples from vaccinated mice or negative control C57BL/6J-A0201 serum (Jackson Laboratory, custom product) serially diluted in 1x ELISA assay buffer (Invitrogen, Cat. #DS98200) were added and plates were incubated overnight at 4°C. Plates were then washed 3 times with 200 μl/well of 1x wash buffer (Invitrogen, Cat. #WB01) and then 100 μl/well of 0.2 μg/ml HRP-conjugated donkey anti-mouse IgG polyclonal antibody (Invitrogen, Cat. #A16017) diluted in 1x ELISA assay buffer was added and plates were incubated for 2 hrs at RT. Plates were then washed 3 times with 200 μl/well of 1x wash buffer and then 100 μl/well of TMB stabilized chromogen (Thermo Scientific, Cat. #SB01) was added. Reactions were allowed to develop for ∼3 minutes and then were stopped by addition of 100 μl/well of ELISA stop solution (Invitrogen, Cat. #SS03). To determine OD_450_ values, plates were read on a Molecular Devices Versamax microplate reader within 60 minutes of addition of stop solution. OD_450_ values were adjusted to account for path length by dividing by the correction factor of 0.596 (corresponding to a volume of 200 μl/well). All buffers were diluted to 1x using UltraPure dH_2_O (Invitrogen, Cat. #10977023). ANOVA analysis was performed in R (v4.5.2) in RStudio (v2025.05.2, Build 517). Raw ELISA data is provided in **Supplementary Data File 2**.

## End Matter

### Author Contributions

JR led the design of the constructs and performed construct generation. SB performed structural predictions. JR and LD performed IVT reactions. LD performed mRNA LNP encapsulations and mRNA LNP QC. TL performed ELISA. CD performed all other *in vitro* work. All *in vivo* procedures were performed by the BC Cancer Investigational Drug Program (IDP). CD and RH wrote the paper.

## Acknowledgments

This work was supported by a Cancer Research UK (C54768/A29062). GlycoNet, supported by the federal government’s Strategic Science Fund, provided funding support for this research (GlycoNet project number RP-10). CD was supported by Canadian graduate scholarships from the Canadian Institutes of Health Research (CIHR).

---

**Figure S1.**
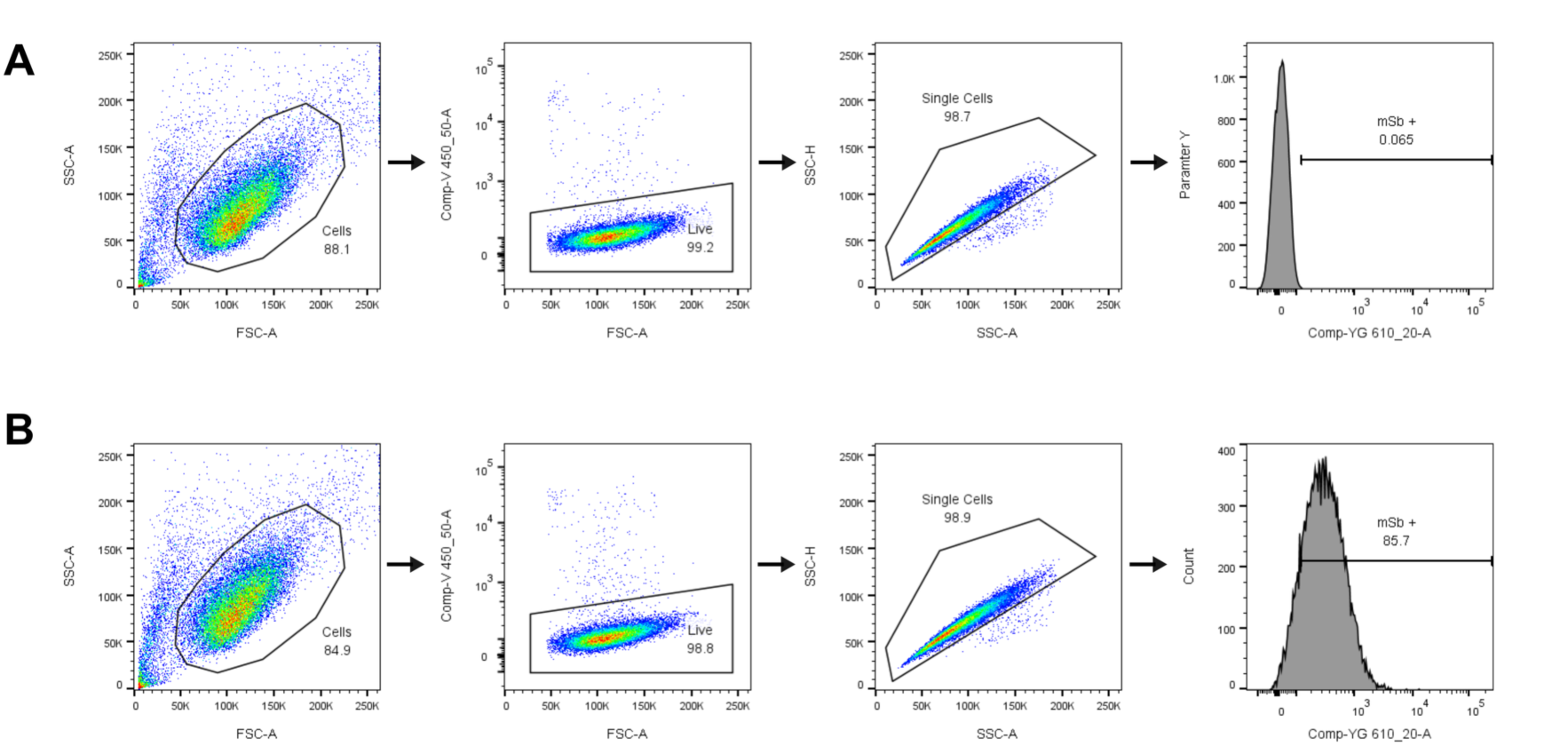
Representative flow cytometry gating scheme for evaluation of secreted monomeric and multimeric mSb-encoding mRNA LNP. Gating for representative negative (A) or positive cell populations (B) related to Figure 2A/B. First, cells are gated based on FSC-A and SSC-A. Then, live cells are gated based on V 450/50A, representing DAPI staining. Then, single cells are gated based on SSC-A and SSC-H. Lastly, cells are evaluated for YG 610/20A signal, representing mSb fluorescence.

**Figure S2.**
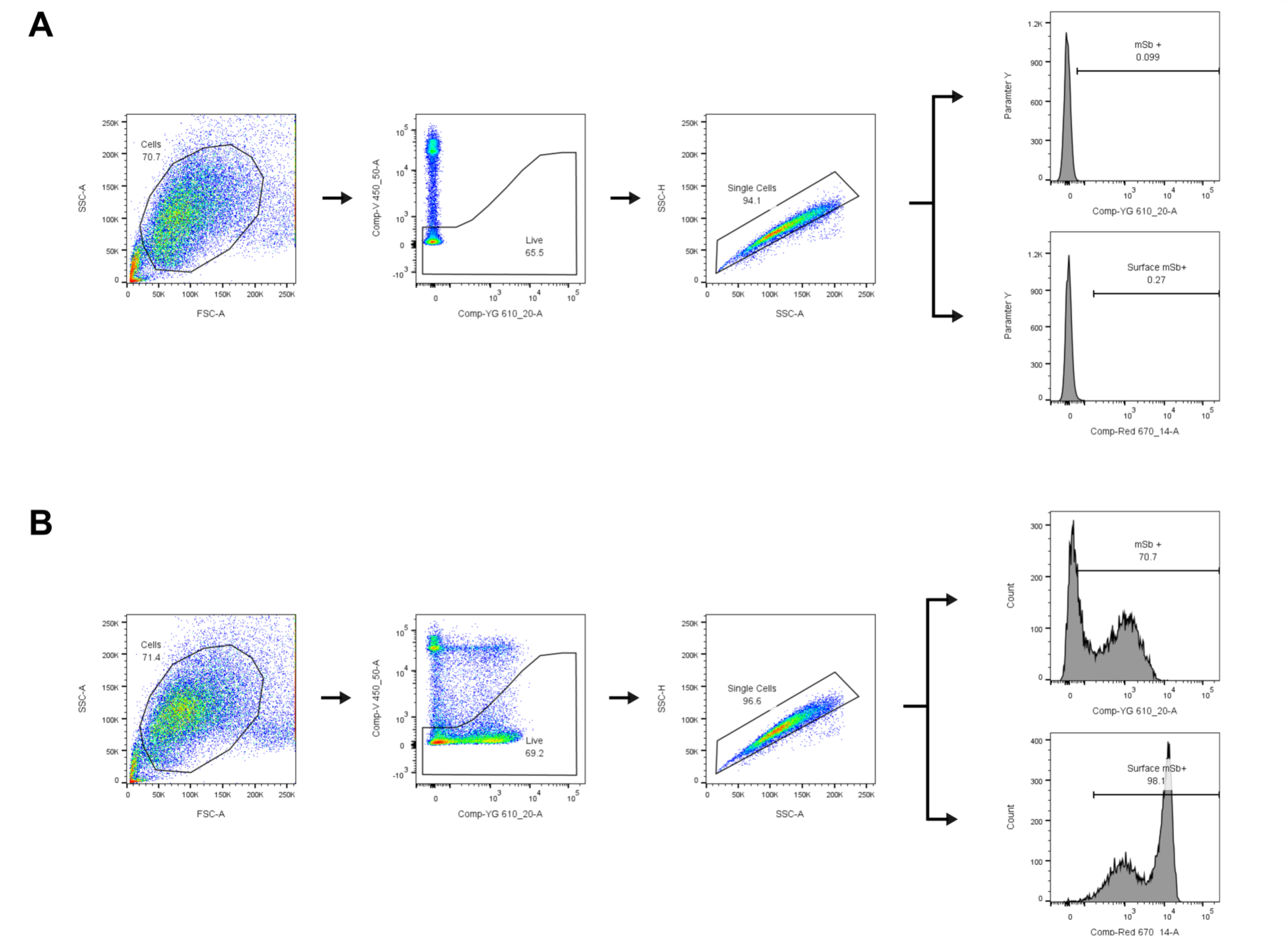
Representative flow cytometry gating scheme for evaluation of surface-displayed mSb-encoding pDNA constructs used to generate mRNA LNP vaccines. Gating for representative negative (A) or positive cell populations (B) related to Figure 3A. First, cells are gated based on FSC-A and SSC-A. Then, live cells are gated based on V 450/50A, representing DAPI staining (note: gating is based on a no viability dye control). Then, single cells are gated based on SSC-A and SSC-H. Lastly, cells are evaluated for YG 610/20A signal and R 670/14A signal, representing mSb fluorescence and mSb surface staining, respectively.

